# Removing Contaminants from Metagenomic Databases

**DOI:** 10.1101/261859

**Authors:** Jennifer Lu, Steven L. Salzberg

## Abstract

Metagenomic sequencing of patient samples is a very promising method for the diagnosis of human infections. Sequencing has the ability to capture all the DNA or RNA from pathogenic organisms in a human sample. However, complete and accurate characterization of the sequence, including identification of any pathogens, depends on the availability and quality of genomes for comparison. Thousands of genomes are now available, and as these numbers grow, the power of metagenomic sequencing for diagnosis should increase. However, recent studies have exposed the presence of contamination in published genomes, which when used for diagnosis increases the risk of falsely identifying the wrong pathogen.

To address this problem, we have developed a bioinformatics system for eliminating contamination as well as low-complexity genomic sequences in the draft genomes of eukaryotic pathogens. We applied this software to identify and remove human, bacterial, archaeal, and viral sequences present in a comprehensive database of all sequenced eukaryotic pathogen genomes. We also removed low-complexity genomic sequences, another source of false positives. Using this pipeline, we have produced a database of “clean” eukaryotic pathogen genomes for use with bioinformatics classification and analysis tools. We demonstrate that when attempting to find eukaryotic pathogens in metagenomic samples, the new database provides better sensitivity than one using the original genomes while offering a dramatic reduction in false positives.

## Introduction and Background

### Next-generation Sequencing in Pathogen Discovery/Diagnosis

Next-generation sequencing (NGS) over the last few years has emerged as a valuable tool for human pathogen discovery and diagnosis. In the case of human pathogen detection, traditional histological, immunological, or molecular techniques are limited and often yield an incorrect or incomplete diagnosis [1]. As sequencing has grown faster and cheaper, clinicians have begun to explore the possibility of replacing older methods with NGS, which provides a fast, specific, and relatively unbiased method of capturing the full spectrum of macro- and microorganisms in any sample.

A growing number of case studies illustrate the potential for NGS in diagnosis. For example, in 2013 Loman et al. conducted a retrospective investigation into the 2011 German outbreak of Shiga-toxigenic *Escherichia coli* (STEC) [2]. In this study, sequencing led to rapid and accurate identification of the bacterial infection in fecal specimens of the infected patients. In 2014, Hasman et al. analyzed 35 urine samples from patients with suspected urinary tract infections, confirming cultured bacterial infections using sequencing of isolated and cultured bacteria [3]. They also successfully identified polymicrobial bacterial infections by directly sequencing the urine samples. Later in 2014, Wilson et al. used next-generation sequencing of cerebrospinal fluid (CSF) to identify and treat a bacterial *Leptospira* infection in a 14-year old patient [4]. In 2016, Salzberg et al. tested the possibilities of detecting pathogens by sequencing brain or spinal cord biopsies from 10 patients presenting with neurologic symptoms with previously unidentified infections [5]. In that study, NGS identified both bacterial and viral infections in selected patients, diagnoses that were confirmed by traditional immunologic techniques.

### Pathogen Discovery Bioinformatics Pipelines

A critical step in using NGS for diagnosis is in the bioinformatics analysis of the millions (or billions) of genomic reads that result from a sequencing experiment. The identification of the sequenced DNA provides the information about the potential pathogenic organisms causing the infection. Because the source of the sample is human tissue, all the studies mentioned above first filtered out human DNA, which is uninformative for pathogen discovery [2–5]. Following this step, the remaining sequencing reads are compared to reference genomic databases, such as RefSeq or the NCBI nt database, using a variety of alignment and classification tools, including BLAST, Bowtie2, MetaPhlAn, and Kraken [6–9].

### Challenges in Relying on Reference Databases

Although databases of sequenced pathogens have grown dramatically larger over the past decade, the dependence on reference databases still presents challenges when used for diagnosis, for at least two reasons: (1) no database contains the full spectrum of all potential human pathogens, and (2) existing reference databases have been found to contain contamination.

Over the past two decades, microbial genome projects have predominantly focused on bacteria and viruses. The GenBank repository [10] contains the majority of genome sequence data submitted by laboratories around the world. As of January 2018, GenBank contained genome entries representing over 54,000 bacterial organisms but only 1,791 fungi and 389 protozoa. The NCBI RefSeq project analyzes and filters the Genbank genome sequences to create a more curated database, which is also widely used [11]. This database also reflects the focus on bacterial and viral genomes, with more than 37,000 bacterial organisms and more than 7,500 viral organisms represented. In contrast, RefSeq contains genomes for only 251 fungi and 82 protozoa (**Table 1**).

**Table 1:** Organisms in Genbank and RefSeq as of January 2018. Total genome counts are based on summaries found at ftp://ftp.ncbi.nlm.nih.gov/genomes/refseq/ and ftp://ftp.ncbi.nlm.nih.gov/genomes/genbank/.

|  | Draft and Complete genomes |  | Complete genomes |  |
| --- | --- | --- | --- | --- |
|  | Genbank | RefSeq | Genbank | RefSeq |
| Bacterial | 54,153 | 37,399 | 5,372 | 5,121 |
| Viral | 10,412 | 7,509 | 10,339 | 7,484 |
| Archaea | 1,861 | 533 | 272 | 235 |
| Fungi | 1,791 | 251 | 26 | 8 |
| Protozoa | 389 | 82 | 3 | 2 |
| Vertebrates | 376 | 238 | 71<sup>a</sup> | 55<sup>a</sup> |
| Plants | 320 | 102 | 3 | 3 |

<sup>a</sup> No complete vertebrate genome exists. The number shown here is the number of organisms with chromosome-level assemblies.

The composition of the reference databases is not representative of the species composition of the natural world, but rather reflects a focus on human pathogens, other species of interest to humans, and the challenges of isolating and sequencing DNA from various species [12]. In many cases, microorganisms are difficult to isolate from their surrounding environments, living among thousands of other species in complex ecosystems [13, 14]. Some microorganisms live in extreme conditions and have gone undiscovered until recently [15]. Other microorganisms are difficult to grow in culture to provide sufficient DNA from which to derive a reference genome. As a result of these constraints, most early research into microorganisms focused on a few easily culturable bacteria [16]. However, studies over the last two decades suggest that culturable bacteria represent only a small fraction of the microorganisms on earth [12, 16–18].

Eukaryotic pathogens comprise an underrepresented group of microorganisms in genomic databases, although they are critically important for the diagnosis of human infections. This group includes a diverse group of species that infect multiple areas in the body; e.g., apicomplexans such as *Plasmodium falciparum*, which causes most cases of human malaria [19], and *Toxoplasma gondii* [20], which may cause neurological defects. Other examples include multiple fungal species belonging to the *Fusarium, Aspergillus, Curvularia*, and *Candida* genera, and amoebae species belonging to the *Acanthamoeba* genus, the latter of which causes a majority of human corneal infections [21, 22]. These are only a small sample of the hundreds of known eukaryotic pathogens of humans.

EuPathDB is a database representing more than 250 eukaryotic microorganisms [23], including both known pathogens and other closely related non-infectious eukaryotic species. Because no eukaryotic pathogen has yet been completely sequenced, this resource comprises primarily draft genomes at varying degrees of completeness, some of which have had little curation since their initial sequencing. However, EuPathDB is more comprehensive than the RefSeq database, containing more than 150 genomes that are absent from the RefSeq protozoa and fungi databases (see **Table 2**).

**Table 2:** EuPathDB Genome Representation in RefSeq. This table shows the number of genomes from the eukaryotic pathogen database that also exist in the Genbank and/or RefSeq databases along with the breakdown of their assembly status within those databases.

| Assembly Status | RefSeq |
| --- | --- |
| Complete Genome | 3 |
| Chromosome | 41 |
| Scaffold | 46 |
| Contig | 5 |
| *Not Represented* | *150* |
| **Total** | **245** |

In recent years, multiple studies revealed contamination in the public genome sequences of many organisms, particularly for draft genomes. In 2011, Longo et al. identified 492 non-primate public databases from NCBI, UCSC, and Ensembl containing human genome sequences [24]. A 2014 study found that portions of the complete genome for *Neisseria gonorrhoeae* TCDC-NG08107 belonged to the cow and sheep genomes [25]. Another study in 2015 identified over 18,000 microbial isolate genome sequences that were contaminated with PhiX174, a bacteriophage used as a control in Illumina sequencing runs [26]. 10% of those 18,000 genomes were published in the literature. In 2016, Kryukov et al. identified 154 non-human genome assemblies containing human sequence fragments that were at least 100bp long [27]. As one example, they discovered that more than 330,000 bp in the reference genome of *Plasmodium gaboni*, a relative of *Plasmodium falciparum*, appears to be contaminating human sequence.

Contamination and incompleteness in reference databases causes bioinformatics analysis of sequencing reads to yield both false positive and false negative results, thereby decreasing the overall reliability of NGS in pathogen diagnostics. False positives, where the wrong pathogen is identified, might in turn lead to inaccurate treatments, with the potential to harm rather than help patients.

In this study, we present a new method for eliminating genomic contamination that can be used on both complete and draft reference genomes. We test our method on a large set of eukaryotic pathogen genomes, yielding a cleaned and filtered eukaryotic pathogen database ready for use in bioinformatics pipelines, including those intended for NGS diagnostics, with decreased false positive and false negative rates.

## Method

The eukaryotic pathogen genomes underwent a multi-step cleaning process to remove both contaminating and non-informative sequences (see Figure 1). Each genome was first split into 100bp overlapping pseudo-reads, with each pseudo-read beginning every 50bp along the genome. The pseudo-reads were then compared to three unique databases, using the Kraken [7] and Bowtie2 [8] classification and alignment programs.

**Figure 1:**
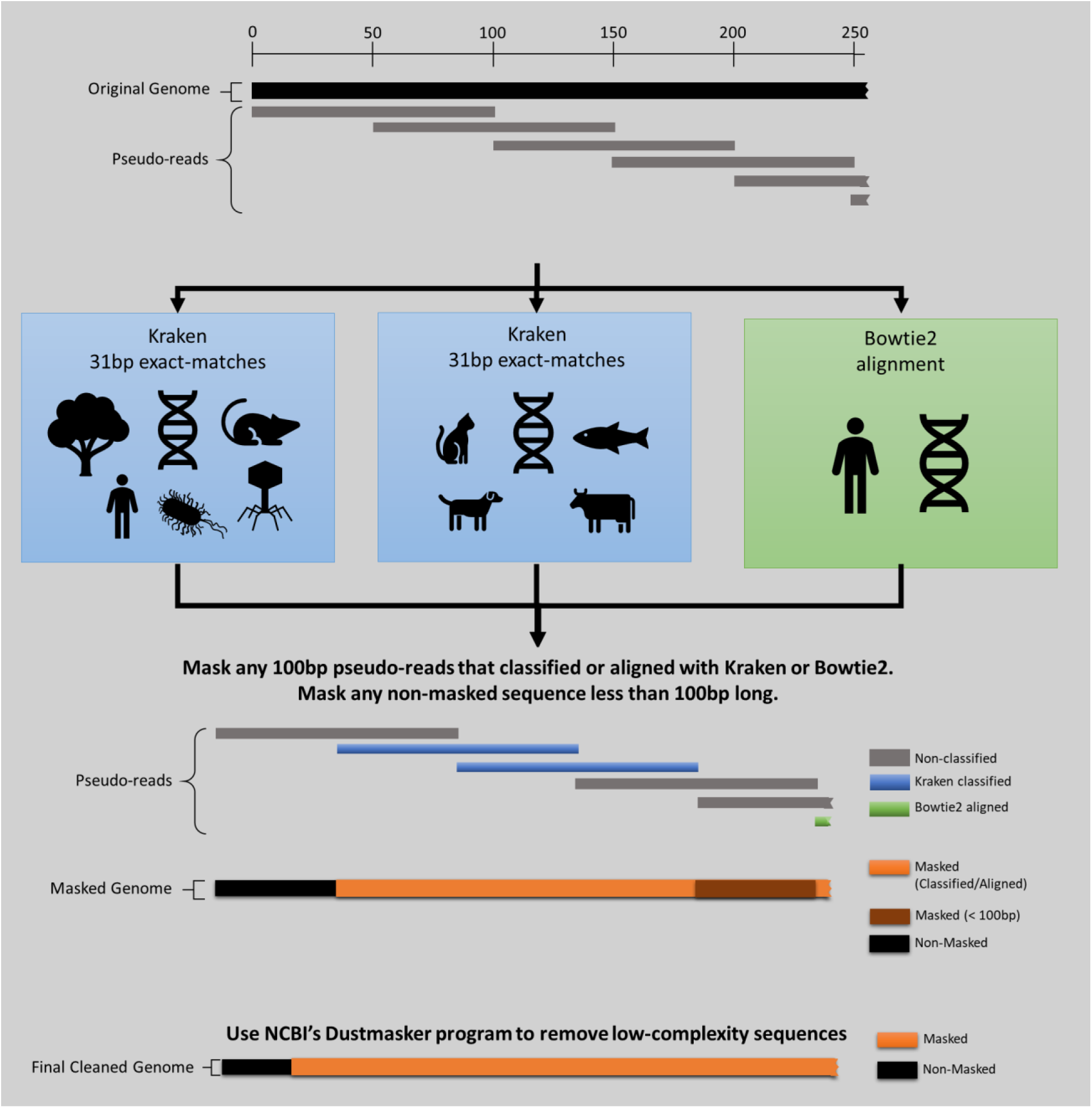
Masking Procedure. A) The original genome is split into 100bp overlapping pseudo-reads. B) The pseudo-reads are then classified using Kraken first against the common contaminating vector sequences and the plant, viral, bacterial, archaeal, human, and mouse RefSeq database. C) The pseudo-reads are also classified using Kraken against non-human and non-mouse vertebrate RefSeq genomes. D) Bowtie2 is then used to align all pseudo-reads against the human genome. E) All pseudo-reads that were classified in the previous steps are masked out of the original genomes. Any remaining non-masked sequence with less than 100p is also masked. F) Finally, Dustmasker is used to mask additional low-complexity sequences.

Kraken labels reads only if they contain an exact 31 base-pair (31-mer) match to any 31-mer in the database sequences [7]. For this process, pseudo-reads were classified with Kraken against two unique Kraken databases. Two separate databases were used to minimize RAM usage. The first Kraken database contains 15,000 genomic sequences from the human, human CHM1, mouse, bacteria, archaea, viral, and plant RefSeq databases as of November 30^th^, 2017. We also included contaminating sequences such as the UniVec database, EmVec database, and phiX174 vector in the first Kraken database. The second Kraken database contains all complete and chromosomal-level assemblies of non-human and non-mouse vertebrate sequences (representing 56 vertebrate species). All pseudo-reads from the eukaryotic pathogen genomes were then classified against both Kraken databases.

Bowtie2 aligns sequencing reads against any reference sequence, allowing for gaps or mismatches [8]. We created a bowtie2 index of GRCh38.p11 and aligned the pseudo-reads against it. Note that even though we include GRCh38.p11 in the Kraken database, which enables Kraken to find human reads, Bowtie2‘s more sensitive alignment algorithm can align some sequences that Kraken will miss.

Any pseudo-read that was classified in these steps represents either a contaminating sequence in the pathogen genome or a low-complexity sequence that matches a distant species only by chance. In either case, these sequences could lead to false positive identifications if they are used for metagenomics analysis. Therefore, we masked any portion of a database genome that corresponded to pseudo-read that was classified or aligned in the previous steps. (Masking can be done in a variety of ways; we simply replaced the sequence with Ns to keep the overall genome length the same.) If, after this initial masking step, we created non-masked sequences that were <100 bp in length, we masked those sequences as well. We then used Dustmasker [28] to mask additional low-complexity sequences (Figure 1).

### Database Availability

EuPathDB-clean is freely available for download at http://ccb.jhu.edu/data/eupathDB/.

### Results and Discussion

We tested our method for eliminating contamination on the draft genomes contained in EuPathDB release 28 [23], which contains 245 genomes categorized into the following sub-databases: AmoebaDB (29 genomes), CryptoDB (11), FungiDB (87), GiardiaDB (6), MicrosporidiaDB (25), PiroplasmaDB (8), PlasmoDB (9), ToxoDB (30), TrichDB (1), and TriTrypDB (39). **Supplementary Table 2** lists all genomes included in EuPathDB, detailing each genome’s filename, sub-database category, genus, species, and full scientific name. **Figure 2** shows how much of each of the 245 genomes was masked in each step of the cleaning procedure and the final lengths of the cleaned pathogen genomes. Full details of the amount of masked sequence for all genomes are listed in **Supplementary Table 2**.

**Figure 2:**
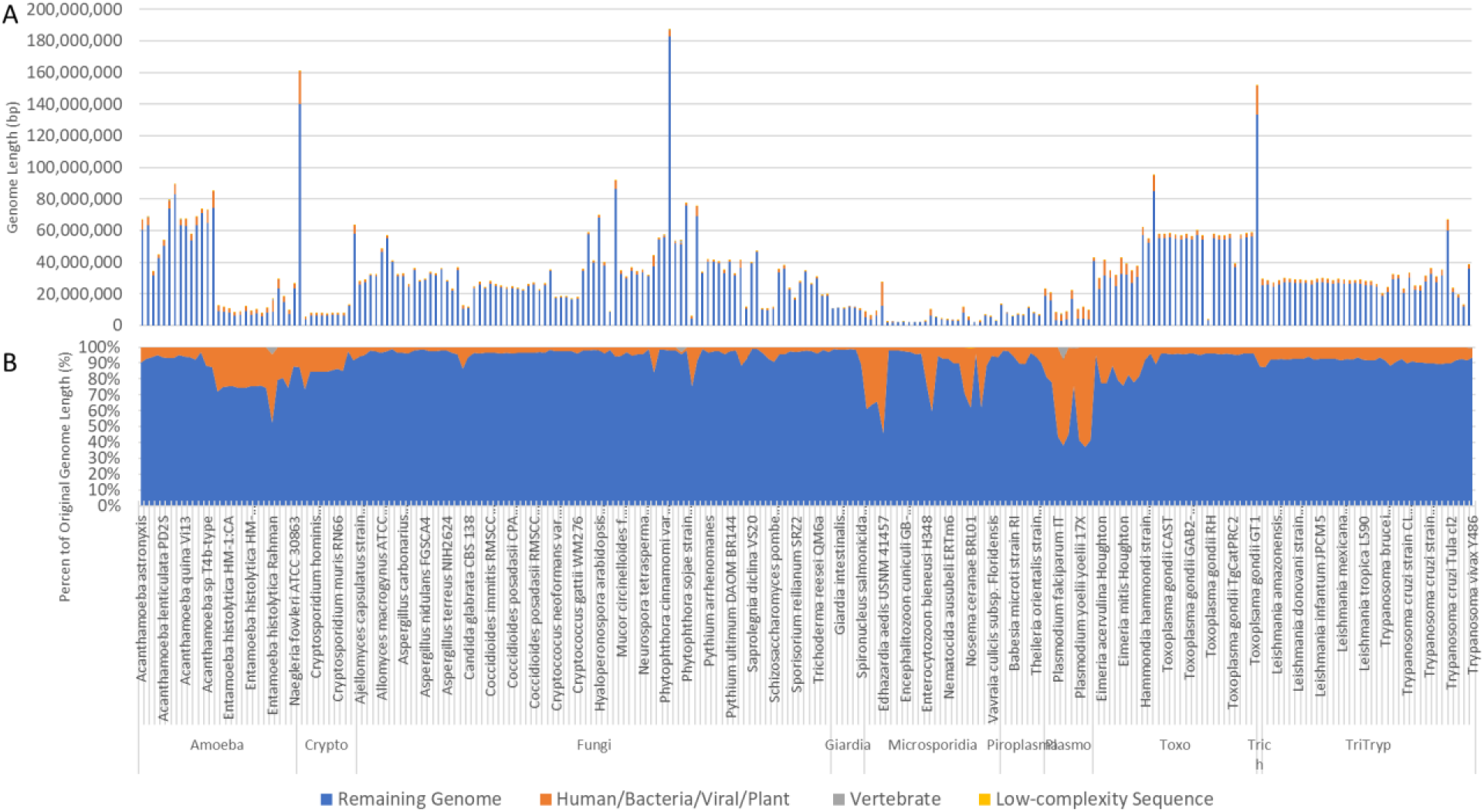
**Masking Results**. Figure 2A displays the genome lengths remaining after each masking step along with the lengths of the genome masked against each database. Low-complexity sequences were masked as a final step as well. Genome lengths are also presented as percentages of the original genome length to show the percent of each genome remaining and the percent masked in each step (Figure 2B).

Genome lengths in EuPathDB ranged from 2Mbp to 186Mbp prior to our cleaning procedure. Post-cleaning genome lengths ranged from 1.7Mbp to 182Mbp, with an average of 11% of each genome identified as contaminating or low-complexity sequences. As Figure 2 illustrates, a few genomes were outliers with over 50% of the genome being masked, but most genomes lost <10% of their length through this process.

In the first masking step, pseudo-reads across all EuPathDB genomes are classified against two Kraken databases containing bacterial, archaeal, viral, human, mouse, vertebrate, and contaminating vector genomes (Figure 1). These classification counts are listed in **Supplementary Table 3.** Reads classified as vertebrates are further broken down into sub-classifications such as fish or bird species. **Figure 3** shows the breakdown of these classifications for the 20 pathogen genomes with the largest numbers of classified pseudo-reads. **Figure 4** shows a similar breakdown focusing specifically on the 20 genomes with the most pseudo-reads labelled as mouse or human.

**Figure 3:**
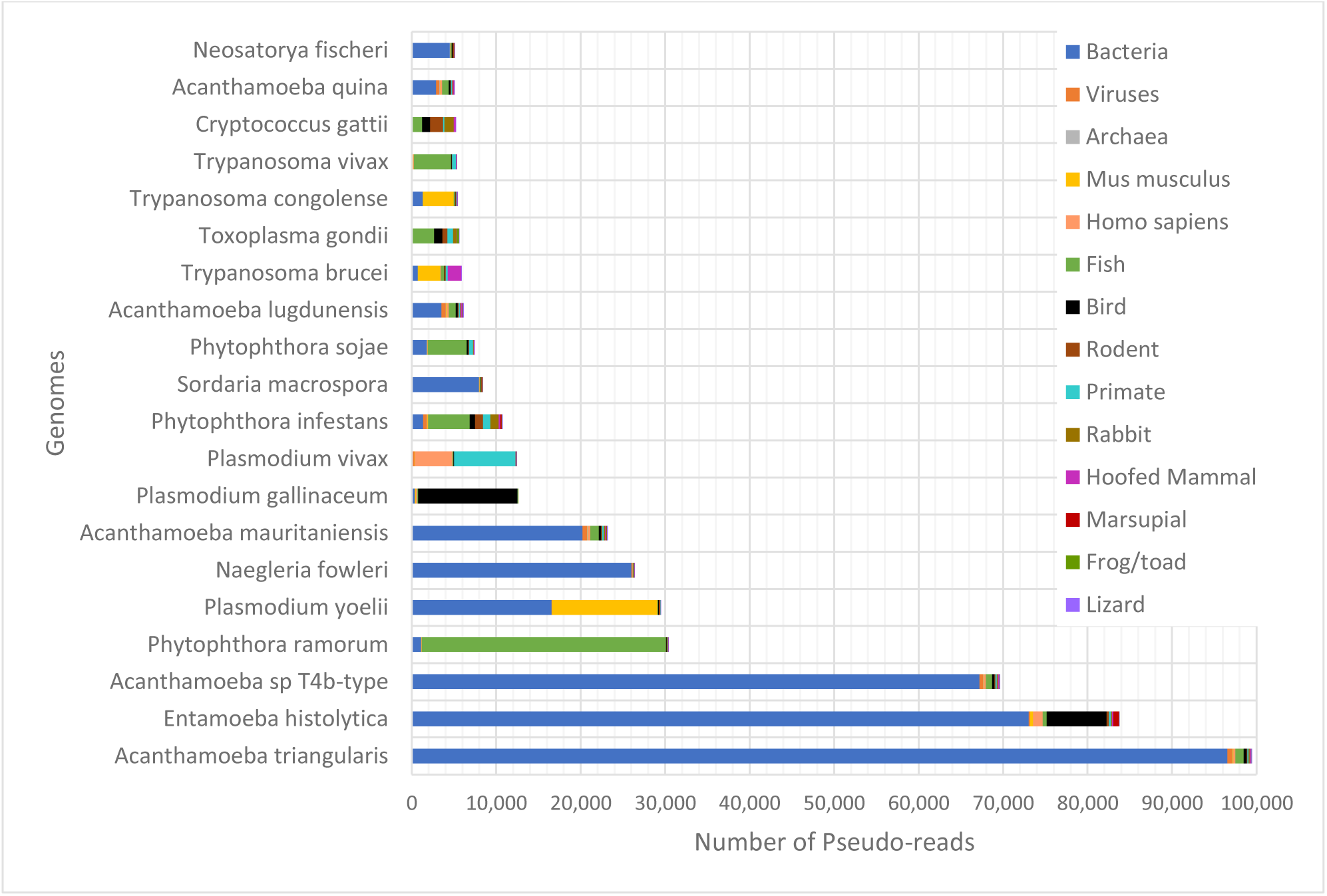
Pseudo-read Kraken classifications. The above plot shows the 20 eukaryotic pathogen genomes with the greatest numbers of pseudo-reads that Kraken identified as matching foreign species when searching against database containing bacteria, viruses, archaea, and a limited set of vertebrate genomes. Vertebrate classifications are grouped by common categories, such as fish, birds, rodents, or primates. Primate and rodent numbers do not include human and mouse, which are counted and shown separately.

**Figure 4:**
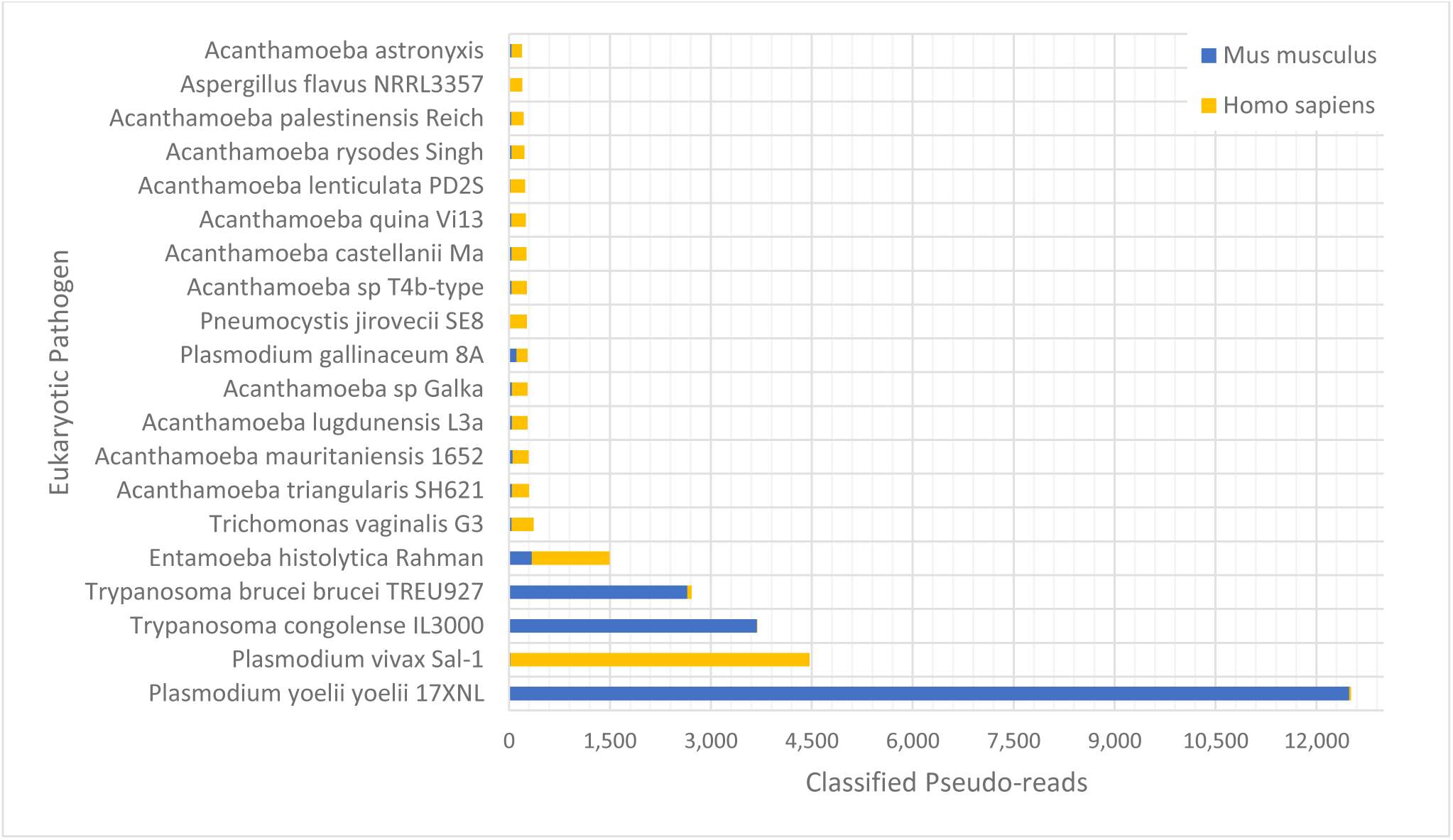
Human/Mouse Classified Pseudo-reads. This plot shows the 20 genomes with the most number of pseudo-reads classified as either human or mouse. Perhaps not surprisingly, the mouse strain of malaria, *P. yoelii*, contains a substantial number of contaminant reads from mouse.

Most genome masking occurred as a result of the first Kraken screen against the database of bacterial, archaeal, viral, human, mouse, and vector genomes. As a result of this step, we masked on average ~10% of each of the EuPathDB genomes. After classifying the remaining pseudo-reads against the vertebrate database, we masked a much smaller amount of sequence, with only 0.1% of each genome matching vertebrate sequences in this step.

The most contaminated eukaryotic pathogen genomes are the three *Plasmodium yoelii* genomes, with approximately 60% of the genomes identified as human/bacterial/viral/archaeal (Figures 3 and 4). The primary sources of contamination in these three genomes were *Methylococcus capsulatus* (16,000 pseudo-reads) and the mouse genome (12,000 pseudo-reads). The genome for *Plasmodium vivax* contained the greatest amount of human contamination, with more than 4,000 pseudo-reads classified as *Homo sapiens.*

### Testing the pathogen database against a set of human cornea samples

To measure the effectiveness of our database cleaning method for NGS diagnosis of human infections, we evaluated a set of 20 human cornea samples recently described by Li et. al 2018 [29] against our EuPathDB-clean. The 20 corneal samples include 4 bacterial infections, 9 eukaryotic pathogen infections, 3 herpes virus infections, and 4 controls. Details about these samples and the true positive pathogens in each sample are listed in **Table 3**, with additional clinical information listed in **Supplementary Table 4.**

For testing, we used 4 Kraken databases: the original EuPathDB, EuPathDB-clean, RefSeq EuPathDB, and a general Microbe Database. The RefSeq EuPathDB contains all protozoal and fungal genomes from the RefSeq database as of December 2017. The Microbe database contains all RefSeq complete bacterial, archaeal, and viral genomes as of December 2017, and it also includes EuPathDB-clean. Genomes contained in each of the above databases are listed in **Supplementary Table 5.**

We first used Bowtie2 to align all corneal sample reads against the human genome reference, GRCh38.p7, and extracted any unaligned reads for each sample (**Table 3**). The non-human reads from each sample were then classified against each database using Kraken. **Supplementary Table 6** contains the full set of species and genus identified in the corneal samples when classified against each database.

**Table 3:** Cornea Sample True Positives. This table summarizes the pathogens present in each of the corneal samples. Metagenomic shotgun sequencing was performed on all samples as described in [29] generating from 20–46 million pairs of 75-bp reads per sample. Sequencing was done in two batches of 10 samples each, where the 10 samples were multiplexed.

| Case # | True Positives | Total 75-bp Paired Reads | Non-Human Aligned Reads |
| --- | --- | --- | --- |
| Case 1 | *Staphylococcus aureus* | 35,947,243 | 8,166,922 |
| Case 2 | *Streptococcus agalactiae* | 42,281,022 | 2,354,821 |
| Case 3 | *Mycobacterium* | 32,321,057 | 1,440,343 |
| Case 4 | *Mycobacterium chelonae* | 31,259,428 | 2,927,088 |
| Case 5 | *Candida* | 22,572,576 | 3,615,840 |
| Case 6 | *Fusarium solani* | 43,187,311 | 3,048,256 |
| Case 7 | *Candida albicans/dubliensis* | 45,410,366 | 1,993,853 |
| Case 8 | *Curvularia* | 42,359,755 | 3,181,901 |
| Case 9 | *Aspergillus flavus* | 46,033,752 | 2,875,199 |
| Case 10 | *Anncaliia algerae* | 20,060,037 | 2,756,229 |
| Case 11 | *Acanthamoeba* | 43,742,352 | 2,880,293 |
| Case 12 | *Acanthamoeba* | 46,648,496 | 3,602,638 |
| Case 13 | *Acanthamoeba* | 44,554,101 | 3,472,961 |
| Case 14 | *Herpes simplex type 1* | 22,460,961 | 1,470,059 |
| Case 15 | *Herpes simplex type 1* | 25,512,845 | 1,411,580 |
| Case 16 | *Herpes simplex type 1* | 23,749,398 | 3,874,558 |
| Case 17 | None | 43,643,461 | 2,637,693 |
| Case 18 | None | 45,824,224 | 2,341,716 |
| Case 19 | None | 25,623,975 | 1,071,939 |
| Case 20 | None | 25,202,226 | 1,823,615 |

Figure 5 summarizes the results when using each of the four databases to identify the pathogens in these samples. The classifications differed greatly depending on the database used, demonstrating the importance of database selection prior to the computational analysis of any NGS sample. However, in the case of diagnostics, the contamination in the raw (unprocessed) genome databases creates false positive signals that overwhelm the true pathogen in of the samples. Classification with the RefSeq EuPathDB yields a similar distribution of microbes for every corneal sample (**Figure 5B**). The resulting read counts suggest that each cornea has a significant presence of *Magnaporthe oryzae*, a pathogen that infects rice plants, and *Toxoplasma gondii* [30]. Similarly, classification against the original EuPathDB presents *Toxoplasma gondii* as one of the primary infections in all but one of the corneal samples (**Figure 5A**). None of the cornea samples had infections by either *Magnaporthe oryzae* or *Toxoplasma gondii* [29],thus both of these classifications are false positives.

**Figure 5:**
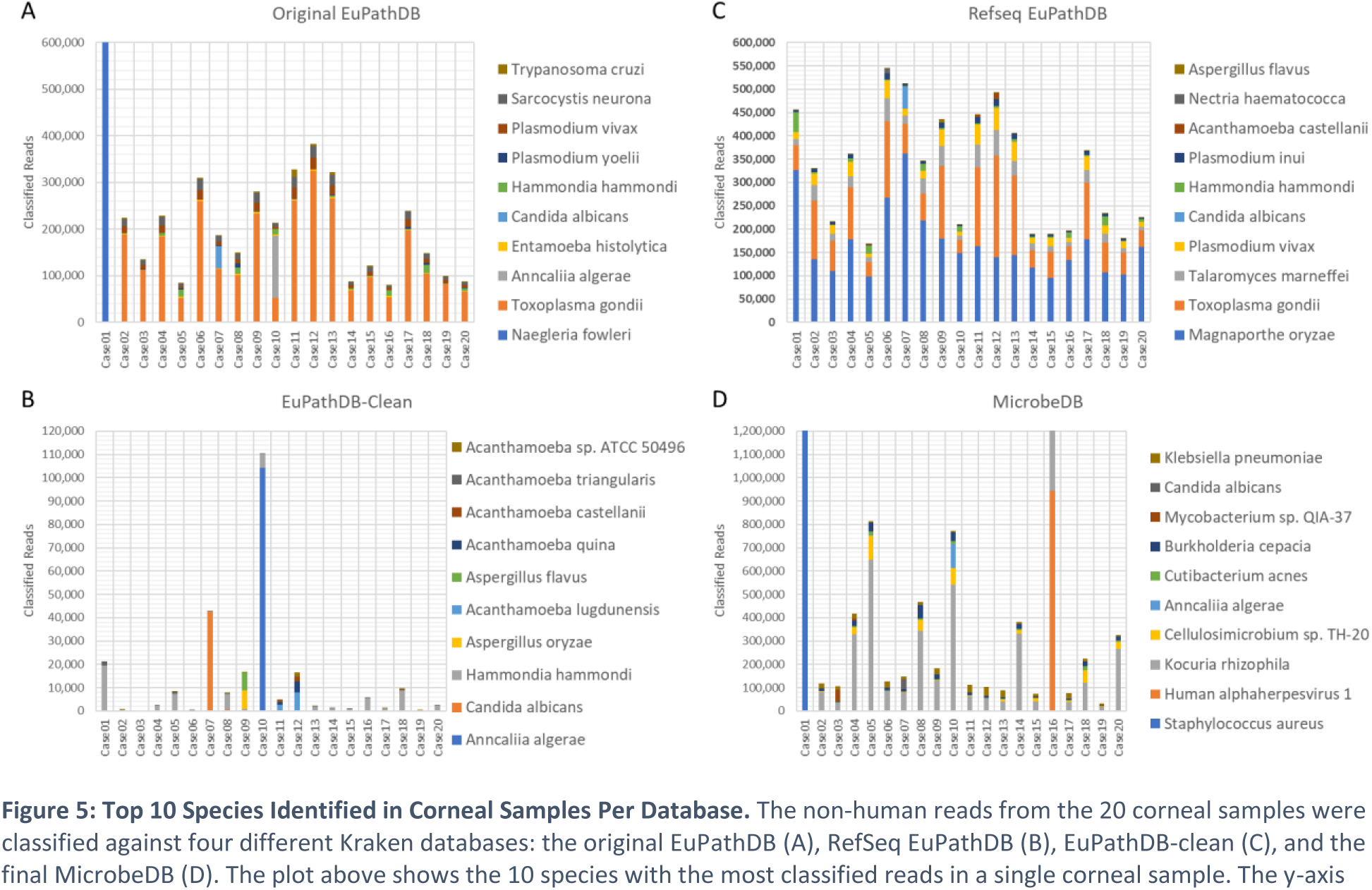
Top 10 Species Identified in Corneal Samples Per Database. The non-human reads from the 20 corneal samples were classified against four different Kraken databases: the original EuPathDB (A), RefSeq EuPathDB (B), EuPathDB-clean (C), and the final MicrobeDB (D). The plot above shows the 10 species with the most classified reads in a single corneal sample. The y-axis for the original EuPathDB and the MicrobeDB are cut off at 600,000 and 1,200,000 reads each to better show the read distributions across all corneal samples.

The contamination removal process masked on average 5% of each *Toxoplasma gondii* genome. For example, the initial *Toxoplasma gondii ME49* genome is ~60 Mb long and the final masked genome is 57 Mb. Fortunately, removing this relatively small proportion of the genome produced a cleaned database with a far better classification profile for the corneal samples. As shown in **Figure 5C**, the correct eukaryotic infections for Cases 7, 9, 10, 11, and 12 are immediately evident with the new database. Instead of hundreds of thousands of reads identified as *Toxoplasma gondii*, the new database shows very high (and correct) read counts for *Anncaliia algerae* in Case 10, *Candida albicans* in Case 7, *Aspergillus* in Case 9, and *Acanthamoeba* in Cases 11 and 12, all true positive infections. With EuPathDB-clean, the maximum number of reads labeled as *Toxoplasma gondii* in any single sample was 18.

After combining EuPathDB-clean with the RefSeq prokaryotes to create MicrobeDB (Figure 5D), we still found a strong signal for the eukaryotic pathogens in their corresponding true positive samples; e.g., the signal from *Anncaliia algerae* in Case 10 in **Figure 5D**. We note that other microbial contamination appears evident when using this database: in particular, *Kocuria rhizophila* appears in every sample, often at high levels. This does not appear to be a database error, as the *K. rhizophila* genome shows now sign of contamination. Instead, the reads from *K. rhizophila* are likely a consequence of physical contamination of the samples at some point in the process.

Another way to look at the data is to examine the read counts for the true positive species only, as shown in Figure 6. Here we show the number of reads in each sample that were assigned to 10 pathogens known to be present in at least one of the samples. With EuPathDB (Figure 6A), at least six samples appear to contain Acanthamoeba, although only samples 11, 12, and 13 came from true Acanthamoeba infections. The RefSeq EuPathDB performed much better, identifying the correct pathogen with a strong signal for Fusarium, Candida, Aspergillus, and 2 cases of Acanthamoeba. EuPathDB-clean had the best signal-to-noise ratio, with strong positive evidence for 5 correct species. RefSeq EuPathDB missed the *Anncaliia algerae* infection because that genome is missing from that database. EuPathDB-clean found Anncaliia, but its signal for Fusarium was weaker (although the signal was still present; see **Supplementary Table 6**). Finally, we note that the genome of Curvularia clavata, the true pathogen in Case 8, has never been sequenced, and therefore none of the methods were able to identify it.

**Figure 6:**
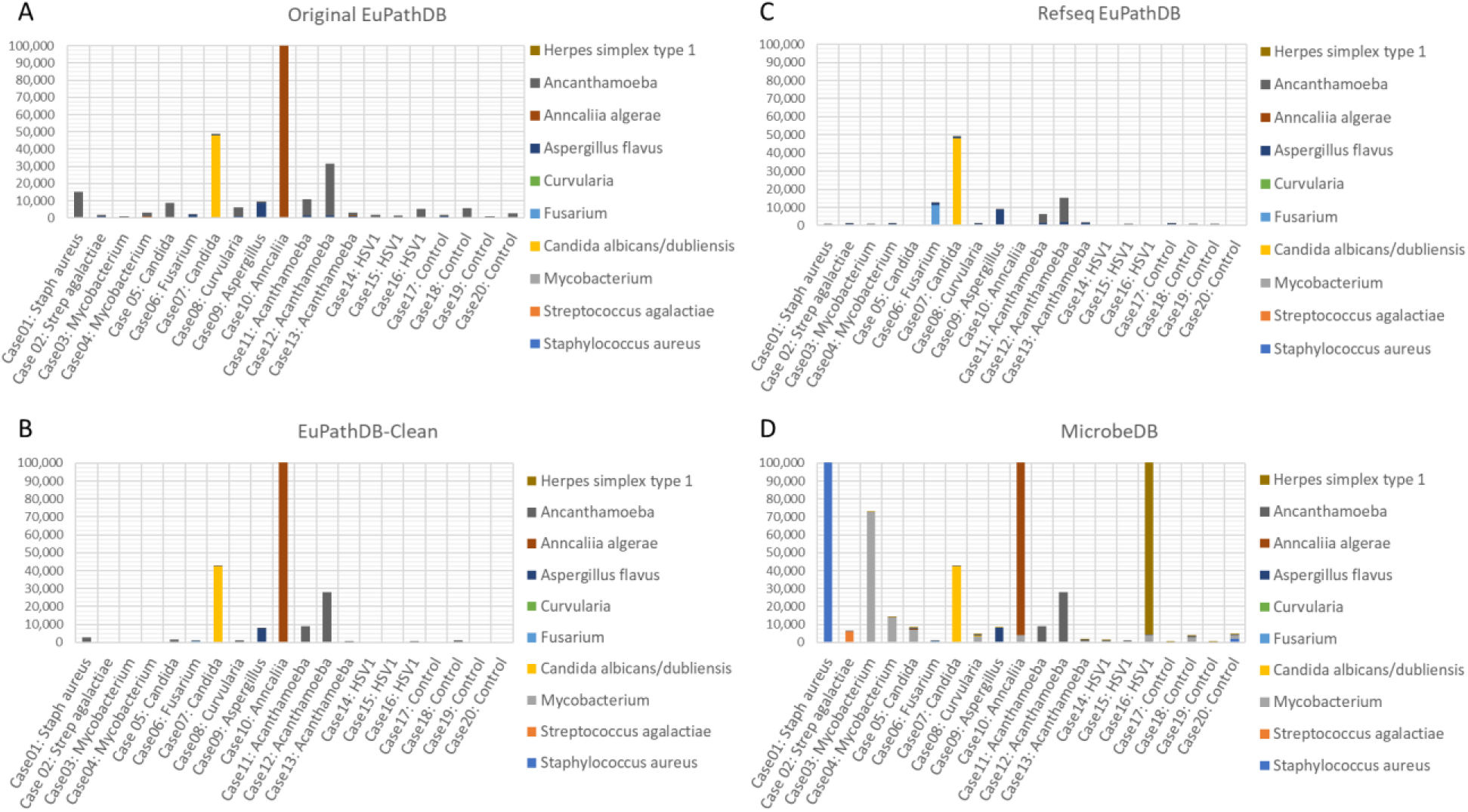
True Positive Species/Genera Identified in Corneal Samples Per Database. The above plot shows only the read counts for the true pathogens present in at least one of the corneal samples.

## Conclusion

In principle, next-generation sequencing can identify all microbial organisms within any sample, making it a potentially a revolutionary method for the diagnosis of human infections. However, this method relies heavily on the computational analysis that compares sequencing reads against reference databases, such as RefSeq and GenBank. Although new genomes are being sequenced daily, the reference databases remain incomplete and, because most new genomes are in draft form, inaccurate. Recent studies have identified contamination in many published genomes, hindering our ability to use them for accurate diagnosis.

We therefore developed a comprehensive contamination removal process, identifying human, vertebrate, bacterial, viral, archaeal, and vector contamination in 245 eukaryotic pathogen draft genomes. By removing contamination and low-complexity sequences, we have created a much cleaner database that minimizes false positives and provides better identification of true positive pathogens in NGS diagnostic samples.

**Supplementary Table #1: Masking Database Composition.** This table lists all genomes used to filter the Eukaryotic Pathogen genomes, including genome accessions, taxonomy IDs, species names, and strain specific names.

**Supplementary Table #2: Eukaryotic Pathogen Database Genomes**. Each genome is listed along with their filename, sub-categories, genus, species and genome lengths ***pre-/post-cleaning.***

**Supplementary Table #3: Pseudo-read counts**. This table lists the number of pseudo-reads for each eukaryotic pathogen that were mistakenly classified as Bacteria, Archaea, Viral, Human, Mouse, or a number of vertebrate categories (i.e. fish, bird, mammal, etc)

**Supplementary Table #4: Cornea Samples Test Case**. Each cornea sample is listed alongside the clinical diagnosis, microbiology test result, and the expected true positive pathogen

**Supplementary Table #5: Testing Database Composition.** This table lists all genomes included in the RefSeq Eukaryotic Pathogen database and the MicrobeDB database.

**Supplementary Table #6: Cornea Sample Classifications.** These tables contain the genus and species classifications assigned by Kraken for the 20 corneal samples when using the original EuPathDB, EuPathDB-clean, the RefSeq EuPathDB, and the MicrobeDB databases.

